## Supplementary Information for "A high-content RNAi screen reveals multiple roles for long noncoding RNAs in cell division"

**The PDF file includes:**

Supplementary tables and movie legends

Sequences of *linc00899* and *C1QTNF1-AS1* vectors (Labomics)

Supplementary figures 1-14

**Supplementary Table Legends**

**Supplementary Table 1.** The list of 2231 lncRNAs in Lincode library.

**Supplementary Table 2.** Raw data and the Z-scores for mitotic progression (mitotic index), chromosome segregation and cytokinesis after depletion of 2231 lncRNAs in screen A and B.

**Supplementary Table 3**. Raw data for the third validation screen (57 lncRNA candidates including Control si (Ambion), GNG12-AS1 (exon 1), Ch-TOG/CKAP5 and ECT2 siRNA).

**Supplementary Table 4**. The list of Stellaris FISH probes against *linc00899*, *TPPP* and *C1QTNF1-AS1.*

**Supplementary Movie Legends**

**Movie S1.** Mitosis in HeLa Kyoto cells transfected with control siRNA (Ambion).

**Movie S2**. Mitosis in HeLa Kyoto cells transfected with *linc00899* siRNA.

**Movie S3**. Mitosis in HeLa Kyoto cells transfected with *C1QTNF1-AS1* siRNA.

**Movie S4**. Mitosis in HeLa Kyoto cells transfected with control LNA A.

**Movie S5**. Mitosis in HeLa Kyoto cells transfected with *linc00899* LNA1.

**Movie S6**. Mitosis in HeLa Kyoto cells transfected with *C1QTNF1-AS1* LNA1.

For all the movies, images were analysed by time-lapse microscopy using Zeiss Axio Observer Z1 microscope. Frames were acquired every 10 min. HeLa Kyoto cells are stably expressing histone H2B-mCheery (red, chromatin marker) and eGFP-$\alpha$-tubulin (green, microtubule marker).

**LncRNA sequence from *Labomics***

**>Linc00899_full sequence**
cggccgcccccgaagcgctgctgtcaccccggccgcgccccccaactttctgcacagtcgcggagctggaagtttcc
ggg
cttcgcggacacgctgggctgggtttcagtcgcggctccgaggttggcaacaaagagggaaagaaggaggaaaa
gcaggc
cggggaggggaggaagagaaccgcgcggaggccgcggcgccgagagccccagaacttccaattctacccaga
agcttttt
tcgtcgtgtttttctcttagacatgatcctctctgaggttggtcctgggcttccatacgtgattcatggaagaggtctca
gccccaagagcccctgagggtactgtccactcccccctggaaacttccagaacctgacgtggggctgaagacataga
ggct
ctgagagttacataattgattctgactttggctgttggtcaacagtgtcataaggtaaaataaggctgttgtagaatctg
ctcagccagggaggggaaaggagtcggcaatcaggtctcctcctgggcacctttgtgaggccagctggcgagagtg
gggg
gtgacactgaggtcccagcagctccaaatgcaggcagagccctgtcctcagagaaggtcacagcctagccaagcc
cagcc
aggtggatgggcccacggaacgcacaggaacctggaacggaggttgaaagcaggaagcacagtctgtgactcc
ccagccc
actctgcattcgaccacttggggcccagaagcttcaggaaaggtgcacaaggtcactgggtcccagtactcccaaca
gga
aggtctggtccagggacagggctcttcccgactccccttagccacacgcaccagaagttctgcagtgcccagtgggc
ata
gcagtccccaagaatgacccagcactgaagctgagccaaagaacttggggagcgagccacacccccctcactccc
tcttcg
cctgctccagacttgccaggcggttgcttttcgtgggaaccgggatgtcctcaccaccctgtccagggcccagccccat
g
tccctggcctgctacagctggaaaaaaaaaagagagatgtttgtttttatttgtttataaaaagaaaagtgttatatata
taacatattatacctcatgaatacatacaattatttgtcaattaacaataaagaaaaatacagcaagcaaaaaagact
ct
cttccacaaaaatagtgttcattacagaaaagtacaaaaaaaaaaaaaaaaaactaagaggatatttagaattaag
aaaa
aactaagagggtatttagaattaaaaaataaaaagaaaacaattacccatgaggtaattactgaatgcatttggttga
aa
gtccttctgctatattttccaaactgtatgtgtatatatgtgcgcatgcatatatgtgtttgtgtgtgtgtacacatatc
tatatgaatggaccatatcgtaagttataaatgcatacatatattcatgtatatataatcattagatcatactataggtt
attttacagtctttttgttgaatacgttgtgagcattttatgtcattattttctacaagatttgaaaaataaagtataaa
taccagttaa

**>Linc00899_scrambled sequence**
ccagctctgaagaacagatgccatgttttaatatctcaagtcggagaaaatatttcgctgggtgcgaagtacttagcat
agatggctccctaggtcattgacatgttctgagacgcttagccaaagtctccatgtgacgtcatatcacaaccacccag
cagcgaagacatgcaaatagatatatgagacgcctcgggtctttcacacaccgtatacctggtcataatctgctccttt
acatcgcgtacttcttgtgtcagggtcttgggggacctttcggaatcatatgtagccggtgtcagacgaaaatgcgtta
tccatcttggcacgtggcgcagtcacctcaagtaagtgcagacaatggttcctatacctttcgtgcacggttgtaataa
tcaaggatggcgttactgttggcaggcaccctatttagccccacgcctcggtctagctggtgcaagctgggtattgtca
acatgctagaaaagtataaagaaccagtgtcacattttcgaagcactcattggaaagagaaacatcttgcgtctcttt
gatatttaatgtaaatggaattgctccatcttacttagaaccgaaagctcagtaaagaaaacactcggggcgcctattg
cgcgtactgaaaaaccgaactgcagtcatcgtggaacagggagaccccaaccgacaataccacaggccaaagg
ccgtac

tgagtcttcgacagttataattgaaaacaatcaactcttcgcgatgataagaaaagttaccccaggattatgtgtggct
ctgagaaagaagaggcggcaagtgctaagaattatatactcggtgacggaagcaacggagtccgtatgcagttac
agct
tcgtcgataaggtttgcgtggatccatggctcccgctaggtgttgttcgcgtccgggcaggcagtgcacaaaaggagc
c
tgaactactgtcagccgaatgactgcggaaaaaatcgttagaaacatcgacataagtgcggcaaatcacagttgca
atc
aggcagtccacgtgtagaatatctcacacaataaggcgcatgcttcccataaatgtattgagaattagcctacatccgg
taggcgtcaacctagtagcgttgatgatcggcagatcacggtgtccgtcgccctcgggtcgtcgtcactacctatacat
acaccacatatacaatggggataaagaaaccagaaaggctatctagaatttgggacatgaacagtgcgttaatcca
aca
cacacagggataagacattccgaacgcgttttcttacgatcgttgataaatgagagggatcgccattcaccagttgag
a
acactgaccccgtcatgagtatcattgaagtacaaaattggaccaagcggacgatttcatgaagtacagtggacccc
cg
atactcgacattccagtttgaaaagaggtatgtccgcgtaaggactttggggcacgacaccaggatatcatatcgcgc
g
gactagttgcaacttacctgcgagggtagcgggcatagacccagagcttctggacggccttcagtggttaggagtac
ac
gggaaaggcaccccagactgaacgcattga

**>C1QTNF1-AS1_full sequence**
gaaggaggaaaggagtgagcatgtcctgctcctgcatgtccctgcttaagctcaggactggcccttccaggccaagg
ac
cccagcatagaccccaggacagggccccaaggatccctggctcatgagagcggcttgctgggctgccccaagag
agcct
gaaggaaacacattgttgagctgagctgacgtcgctgtttcttccagactgctctctaaagtgggcagggtagcgacc
g
gccggctccgatggtgacgtcccactgccaaggggtgggagtggggagagtctccacagagcttcggagaagctg
ctaa
gatggaaaagtggaaacttggcagacagatccagcctccctggccactggcccatgctcgtggctcctggatggcg
ctg
ccacgttctgagcagcttgggacaggtggagatcaggactggcagctgcaaggacacaccagagccacagaaac
taaag
agaatttccaaaaggagtctatggtgaagtctctgaggatgcaaagaagacaaggagaatgaaaatccaatgaaa
gcct
gattgtatttgttgaccttaaggaaagtgattttatggtacagcctctctggaagggagggtgtgttcgctcacagaat
gcaaatacccctttgacccccctaatctttcttctaggagtttctcctacagataaacttagaagggtgctcaaataagta
agttcaaggatatcctctgaagcattgccgtagtataaaaaaagcacagataccctcaaaggacatcattaggggcc
tg
gtaaaataaattccacacagtggaacaccgtgtagctctttagagaataaacagctctctatatgtgatctggaa

**>C1QTNF1-AS1_scrambled sequence**
gtggcggatgttggcttagaatcaggattgtgcagatattgaaccttgttgcctcccaccctgcacagcttgatgaccc
ctaaaagtagcagagcttcacaagtactattgggcaatcgctccggcgatggtgaaaaaacctactcttcaacacggt
c
gagcttggtctaccctcttcagcgaacattacaagaaacagaactcataggcatcaaatgtttgaggaccatggagg
ga
tatgcctagacgcgcggtagcaggcaagatcagggtttaacatggcctcttgaaacttagaggtacagccgccgcgc
gt

ccgcggtagtgcactgcggaaaccgcataatatcgtgaggtaggtgggcgaagtggtcttggtgcagcttgaccttaa
g
tgttaggacccatgtagatagcaaaagcagggcggcggcgtcataaagggttcagcgcacccatgcgaggaagtt
agga
gagaaccaaggacaatcgggccatcaggatgacaggcagaaacgaagtacacacagaagcgatccgtttattaa
cgcaa
cattggcggctatcaacacattcgtcctattgtgtaggttgcatggtaattctgtcatataaggcgtgtccccaacaca
ataaaggttcgccgctaagctcgccaaatccgatcaggcaacaaagtcggatctatactagtctaaagaggcggag
agg
ctagagcttcaatccgccaccgcgaccaattctaagttaggttagtgtcgactgtgaaagcaaagaccggcctcaac
gg
gtagtgagcggggatggactggagtcctaagcacacgctgttgcgctagagaatctaggactcggttactcatcc

**List of primer sequences for qPCR.**

| Expression primers | Forward primer (5'to3') | Reverse primer (5'to3') |
|---|---|---|
| *GAPDH* | CAACAGCCTCAAGATCATCAG | ATGGACTGTGGTCATGAGTC |
| *RPS18* | ATCCCTGAAAAGTTCCAGCA | CCCTCTTGGTGAGGTCAATG |
| *ACTB (β-actin)* | GTTACACCCTTTCTTGACAAA | GTCACCTTCACCGTTCCAGTT |
| *NORAD* | AGCGAAGTCCCGAACGACGA | TGGGCATTTCCAACGGGCCAA |
| *MALAT1* | GACGGAGGTTGAGATGAAGC | ATTCGGGGCTCTGTAGTCCT |
| *U1* | ATACTTACCTGGCAGGGGAG | CAGGGGGAAAGCGCGAACGCA |
| *Linc00899/LOC100271722* (exon 1-2) | gagagccccagaacttccaa | agcctctatgtcttcagccc |
| *Linc00899/LOC100271722* (exon 2-4) | ctgagggtactgtccactcc | ccctggctgagcagattcta |
| *C1QTNF1-AS1/LOC100507410* (exon 1-2) | agctgacgtcgctgtttctt | tccgaagctctgtggagact |
| *C1QTNF1-AS1/LOC100507410* (exon 2-3) | ggacaggtggagatcaggac | tctccttgtcttctttgcatcc |
| *DUBR/LINC00883* | / | QT02433235 |

| | | |
|---|---|---|
| *(QIAGEN)* | | |
| *PP7080/* LOC25845 *(QIAGEN)* | / | QT01159095 |
| *LOC100506835/linc00840 (exon 1-2)* | ttgcgcacgaattttacac | gtggaccttgtctgcatcct |
| *LOC729970 (exon 2-3)* | gaaccacaaagctgaacctca | agttctgagagcacaagggg |
| *TPPP (QIAGEN)* | / | QT00007693 |
| *CCNB1* | TTTGCACTTCCTTCGGAGAGC | AAGGAGGAAAGTGCACCATG TC |

**List of CHART probes (ordered from ATDBio Ltd, School of Chemistry, Southampton, UK).**

| CHART probes | Probe (5'to3') |
|---|---|
| *Linc00899*_CHARTprobe1_exon 1 | cgcgactgtgcagaaagttggggg/iSp18/3BioTEG |
| *Linc00899*_CHARTprobe14_exon 1 | cgaagcccggaaacttccagctcc/iSp18/3BioTEG |
| *Linc00899*_CHARTprobe9_exon 4 | ccccaagttctttggctcagcttc/iSp18/3BioTEG |
| *Linc00899*_CHARTprobe10_exon 4 | tggacagggtggtgaggacatccc/iSp18/3BioTEG |
| *Linc00899*_CHARTprobe10_exon 4_ sense (negative control) | gggatgtcctcaccaccctgtcca/iSp18/3BioTEG |
| *Linc00899*_CHARTprobe14_exon 1_ sense (negative control) | ggagctggaagtttccgggcttcg/iSp18/3BioTEG |

**List of CHART primers.**

| CHART primers | Forward primer (5'to3') | Reverse primer (5'to3') |
|---|---|---|
| *Linc00899*_exon 4 | agtccccaagaatgacccag | gacagggtggtgaggacatc |
| *Linc00899*_exon 1 | ccccaactttctgcacagtc | ccctctttgttgccaacctc |
| *5.8 S* (West et al., 2014) | GGTGGATCACTCGGCTCGT | GCAAGTGCGTTCGAAGTGTC |

**List of antibodies.**

| Antibody | Usage | Dilution | Catalog number |
|---|---|---|---|
| TPPP | Western blot | 1-1000 | NBP2-34031, Novus |
| β-tubulin | Western blot | 1-2000 | T019, Sigma |
| p150 | Western blot | 1-2000 | 610473, BD Transduction Laboratories |
| Cyclin B1 | Western blot | 1-1000 | 12231S, Cell signalling |
| Acetylated α-tubulin | IF | 1-500 | T6793, Sigma |
| EB1 | IF | 1-300 | CRUK Hybridoma bank *(Zyss et al., 2011, JCB)* |
| α-tubulin | Imaging screen, IF | 1-1000 | TUB9026, Sigma |
| Phospho Histone H3 (Serine 10) | Imaging screen, IF<br>Western blot | 1-2000<br>1-1000 | 06-570, Millipore |
| α-tubulin (rat) | Imaging screen, IF | 1-500 | MCA78G, AbD Serotec |
| γ-tubulin | Imaging screen, IF | 1-1000 | GTU88, Sigma |
| CDK5RAP2/CEP215 | Imaging screen | 1-500 | (Barr et al., 2010, JCB) |
| H3K4me3 | CUT&RUN | 1-100 | 05-1339, Millipore (LOT2780484) |
| H3K36me3 | CUT&RUN | 1-100 | 61101, Active Motif (LOT32412003) |
| H3K27me3 | CUT&RUN | 1-100 | 9733S, C36B11, Cell Signaling (LOT8) |
| H3K27ac | CUT&RUN | 1-100 | Ab4729, Abcam (LOT GR3187598-1) |

| | | | |
|---|---|---|---|
| Goat anti-rabbit IgG secondary antibody | CUT&RUN | 1-100 | Ab97047, Abcam (LOT GR254157-8) |
| Rabbit anti-mouse IgG secondary antibody | CUT&RUN | 1-100 | A27022, Thermo Fisher Scientific (LOT RG240909) |
| Alexa Fluor® 555 goat anti-rat | Imaging screen, IF | 1-1000 | A-21434, Thermo Fisher Scientific |
| Alexa Fluor® 488 donkey anti-rabbit | Imaging screen, IF | 1-1000 | A-21206, Thermo Fisher Scientific |
| Alexa Fluor® 647 donkey anti-mouse | Imaging screen, IF | 1-1000 | A-31571, Thermo Fisher Scientific |
| Amersham ECL anti-mouse IgG, HRP antibody | Western blot | 1-2000 | NXA931V, GE Healthcare |
| Amersham ECL anti-rabbit IgG, HRP antibody | Western blot | 1-2000 | NA934V, GE Healthcare |
| Alexa Fluor 568 Phalloidin | Imaging screen | 1-500 | A12380, Thermo Fisher Scientific |

**List of siRNA sequences.**

| siRNA | Sequence (antisense) | Product number |
|---|---|---|
| siGENOME Non-Targeting siRNA Pool #2 (GE Dharmacon) | / | 001206- 14- 20 |
| Negative control siRNA #1 (Thermo Fischer Scientific, Silencer select, Ambion) | / | 4390084 |
| *GNG12-AS1 exon 1 siRNA #1 (Life Technologies, Silencer select) #S59962* | AUUCUUGUUCACGUCGCC G | 4392421 |
| *Lincode Human Linc00899 SMART pool, GE Dharmacon* | CAGAGAAGGTCACAGCCT A<br>GGGGAAAGGAGTCGGCAA T<br>GGACCATATCGTAAGTTAT<br>GACTTTGGCTGTTGGTCAA | R-189507-00-0005 |
| *Lincode Human C1QTNF1-AS1 SMART pool, GE Dharmacon* | AGATAAACTTAGAAGGGTG<br>ACUAGAGGCTGCAGCGGC A<br>TCTATATGTGATCTGGAAA | R-189493-00-0005 |

| | | |
|---|---|---|
| | TCATTAGGGGCCTGGTAAA | |
| *Lincode Human Linc00883/DUBR SMART pool, GE Dharmacon* | / | R-023569-00-0005 |
| *Lincode Human PP7080 SMART pool, GE Dharmacon* | / | R-028371-00-0005 |
| *Lincode Human Linc00840 SMART pool, Dharmacon* | / | R-189275-00-0005 |
| *Lincode Human LOC729970 SMART pool, Dharmacon* | / | R-182791-00-0005 |
| *Lincode Human* NORAD *SMART pool, Dharmacon* | / | R-038095-00-0005 |
| *TPPP* SMARTpool: ON-TARGETplus (GE Dharmacon) | / | L-019695-01-0005 |
| *Ch-TOG/CKAP5* SMARTpool: ON-TARGETplus (GE Dharmacon | / | L-006847-00-0005 |

| | | |
|---|---|---|
| *ECT2* SMARTpool: ON-TARGETplus (GE Dharmacon | / | L-006450-00-0005 |

**List of LNA Gapmer sequences (ordered from Exiqon).**

| LNA | Sequence (5' to 3') | Product number |
|---|---|---|
| Negative control LNA A | AACACGTCTATACGC | 300611-00 |
| Negative control LNA B | GCTCCCTTCAATCCAA | 300615-00 |
| *Linc00899 LNA gapmer_1 (exon 1)* | cgcgactgtgcagaa | 300603-00<br>Design ID:606267-2 |
| *Linc00899 LNA gapmer_2 (exon 4)* | acttctggtgcgtgtg | 300603-00<br>Design ID:380384-2 |
| *Linc00899 LNA gapmer_3 (exon 1)* | cgactgtgcagaaagt | 339511<br>LG00204910-DDA |
| *Linc00899 LNA gapmer_1 (intron 1) #4916* | ctggtgaggaagaaca | 339511<br>LG00204916-DDA |
| *Linc00899 LNA gapmer_1 (intron 2) #4909* | tagtgtatgatagaac | 339511<br>LG00204909-DDA |
| *Linc00899 LNA gapmer_1 (intron 3) #4912* | acatagaatctggaat | 339511<br>LG00204912-DDA |
| *C1QTNF1-AS1, LNA gapmer_1 (exon 3)* | tacggcaatgcttcag | 300603-00<br>Design ID:591242-1 |
| *C1QTNF1-AS1, LNA gapmer_2 (exon 2)* | gctcagctcaacaatg | 300603-00<br>Design ID:591242-2 |

**List of CRISPR pAS guide RNA sequences and single stranded DNA donors (SSODN) containing SV40 polyA sequence (*in italic)*.**

| Targeted lncRNA | Guide-ID | Guide sequence |
|---|---|---|
| C1QTNF1-AS1_guide 70 Forward (5'-3') Cloned into Addgene vector #48138 | Guide 70 (112bp from TSS**)** | caccGTCCCTGCTTAAGCTCAGGAC |
| C1QTNF1-AS1_guide 70 Reverse (5'-3') Cloned into Addgene vector #48138 | Guide 70 (112bp from TSS) | aaacGTCCTGAGCTTAAGCAGGGAC |
| SSODN_ C1QTNF1-AS1_ guide 70 (3'-5') | Guide 70 | GCAAGCCGCTCTCATGAGCCAGGGATCCTTG GGGCCCTGTCCTGGGGTCTATGCTGGGGTCC TTGGCCTGGAAGGG*cacacaaaaaaccaacacacag atctaatgaaaataaagatcttttatt*GTCCTGAGCTTAAG CAGGGACATGCAGGAGCAGGACATGCTCACT CCTTTCCTCCTTCCCGAGCCCCGCAGTGAG |

**List of CRISPRa guide RNA sequences cloned into Addgene vector #60955.**

| Targeted lncRNA | Guide-ID | Guide sequence |
|---|---|---|
| *Linc00899* | Guide 1 (-129bp from TSS) | AGATCGTCCCGAGCGCGCCG |
| *Linc00899* | Guide 2 (+20bp from TSS) | GGGTGACAGCAGCGCTTCGG |
| Negative control | sgRNA 2 | GTGCGATGGGGGGGTGGG TAGC |

**List of PCR primers used for subcloning of *linc00899* (full and scrambled sequence obtained from Labomics) into the lincXpress vector with the Gateway system.**

| Targeted lncRNA | Forward primer (5'to3') | Reverse primer (5'to3') |
|---|---|---|
| *Linc00899 full* | GGGGACAAGTTTGTACAAAAAAGCAGGCTcggccgcccccgaagcgctgctgtc | GGGGACCACTTTGTACAAGAAAGCTGGGTttaactggtatttatactttatttt |
| *Linc00899 scr* | GGGGACAAGTTTGTACAAAAAAGCAGGCTccagctctgaagaacagatgccatg | GGGGACCACTTTGTACAAGAAAGCTGGGTtcaatgcgttcagtctggggtgcct |

**List of PCR primers for CRISPR pAS insertion.**

| Targeted lncRNA | Forward primer (5'to3') | Reverse primer (5'to3') | PCR product size |
|---|---|---|---|
| C1QTNF1-AS1_Guide 70 | tcccttcctgctcacaccgc | aggaccctgccatctccctga | Wild type=358bp, pAS inserted=404bp, cleaved PCR product after BglII should give rise to 2 bends, 149bp and 237bp. |

| | | | |
|---|---|---|---|
| Forward primer for Addgene vector #48138 | CATGATTCCTTCATATTTGCATATAGC | | (used for Sanger sequencing) |
| mU6 | GAGATCCAGTTTGGTTAGTACCGGG | | (used for Sanger sequencing of CRISPRa clones) |

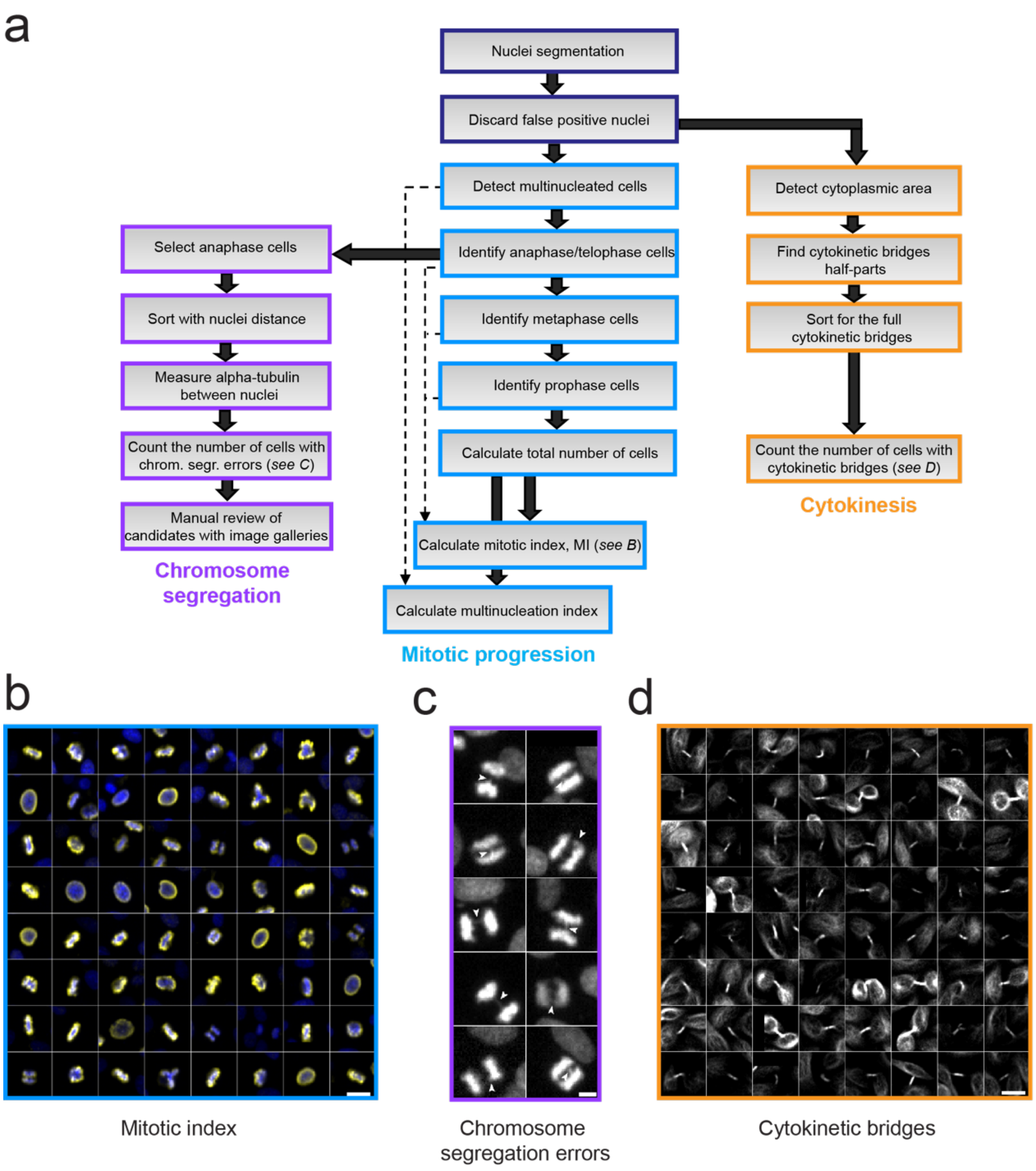


**Supplementary Fig. 1. Workflow to identify new lncRNAs in regulation of cell division.**

***a.*** Schematic of the workflow for automated cell segmentation and quantification of cellular features after depletion of each lncRNA in the Lincode library. This includes the number of mitotic cells (i.e., the mitotic index), the number of chromosome segregation defects in anaphase cells and the number of cytokinesis defects.

***b.*** Gallery of representative images of mitotic cells positive for PHH3 staining (yellow). DNA is stained with Hoechst and is shown in blue. Scale bar, 20 µm.

***c.*** Gallery of representative images of cells with chromosome segregation defects. Arrows mark chromosome segregation errors. Scale bar, 20 µm.

***d.*** Gallery of representative images of cells with cytokinetic bridges. Scale bar, 20 µm.

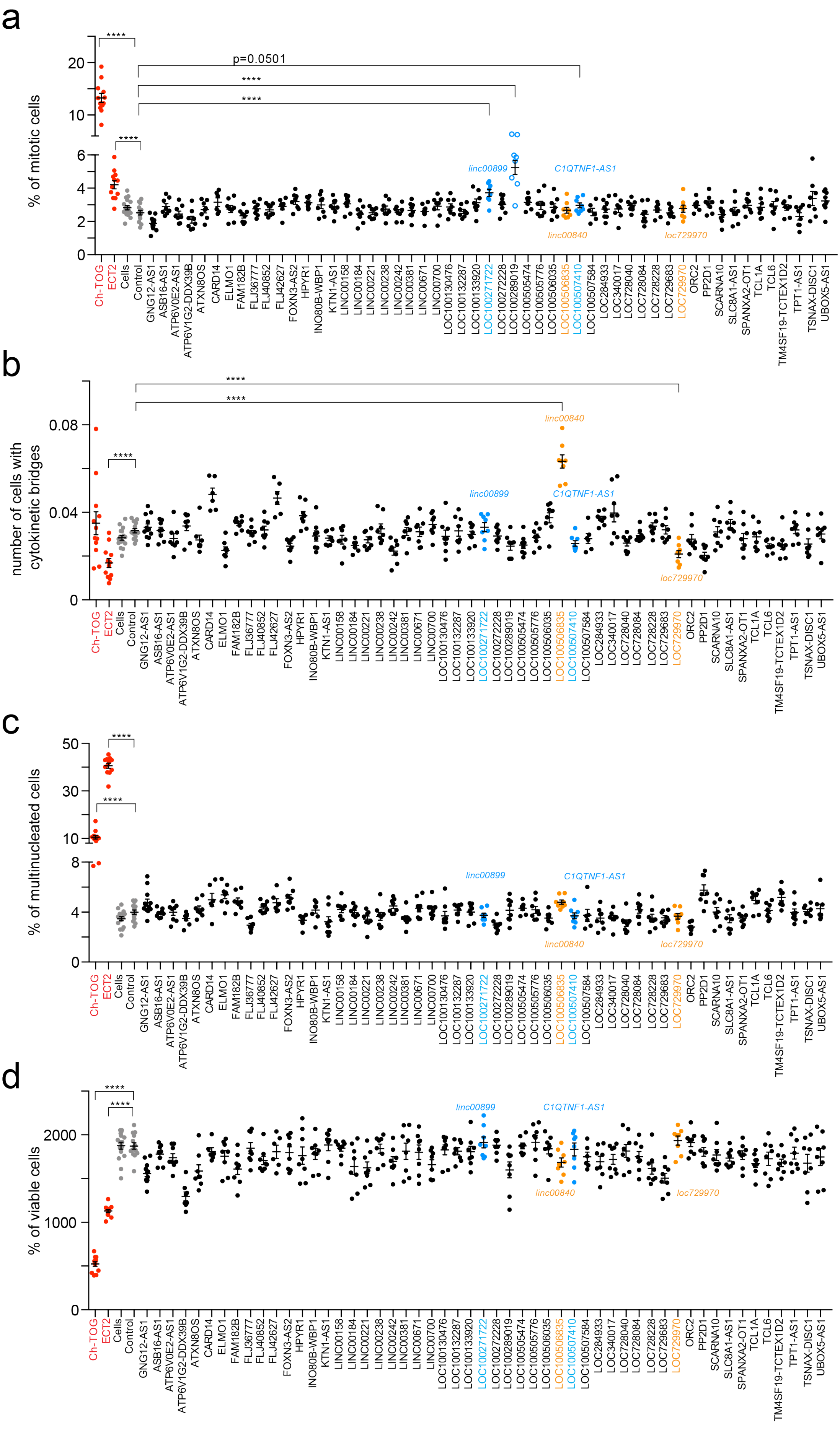
a
% of mitotic cells
20
10
6
4
2
0
****
p=0.0501
****
****
****
linc00899
C1QTNF1-AS1
linc00840
loc729970
Ch-TOG
ECT2
Cells
Control
GNG12-AS1
ASB16-AS1
ATP6V0E2-AS1
ATP6V1G2-DDX39B
ATXN8OS
CARD14
ELMO1
FAM182B
FLJ36777
FLJ40852
FLJ42627
FOXN3-AS2
HPYR1
INO80B-WBP1
KTN1-AS1
LINC00158
LINC00184
LINC00221
LINC00238
LINC00242
LINC00381
LINC00671
LINC00700
LOC100130476
LOC100132287
LOC100133920
LOC100271722
LOC100272228
LOC100289019
LOC100505474
LOC100505776
LOC100506035
LOC100506835
LOC100507410
LOC100507584
LOC284933
LOC340017
LOC728040
LOC728084
LOC728228
LOC729683
LOC729970
ORC2
PP2D1
SCARNA10
SLC8A1-AS1
SPANXA2-OT1
TCL1A
TCL6
TM4SF19-TCTEX1D2
TPT1-AS1
TSNAX-DISC1
UBOX5-AS1
b
number of cells with cytokinetic bridges
0.08
0.04
0
****
****
****
linc00840
linc00899
C1QTNF1-AS1
loc729970
c
% of multinucleated cells
50
30
10
8
4
0
****
****
linc00899
C1QTNF1-AS1
linc00840
loc729970
d
% of viable cells
2000
1000
0
****
****
linc00899
C1QTNF1-AS1
linc00840
loc729970

**Supplementary Fig 2. Validation screen for lncRNAs involved in mitotic progression and cytokinesis.**

Top 25 lncRNAs from screens A and B were selected and the validation screen was performed in HeLa cells stained with the same antibodies as used in screen B (PHH3, $\alpha$-tubulin and $\gamma$-tubulin). Two biological replicates were performed with four technical replicates. The number of mitotic cells (mitotic index, ***a***), the number of cells with cytokinetic bridges (***b***), the number of multinucleated cells (***c***) and the number of viable cells (***d)*** were calculated. *Ch-TOG* and *ECT2* siRNAs (both as red circles) were used as positive controls for mitotic index and multinucleation, respectively. Depletion of *ECT2* led to an increase in multinucleated cells, while depletion of *Ch-TOG* led to an increase in mitotic index. Cells treated with negative control siRNA (Control, Ambion) and cells alone are indicated as grey circles. Top lncRNA candidates involved in mitotic progression and cytokinesis are indicated as blue and orange circles, respectively.

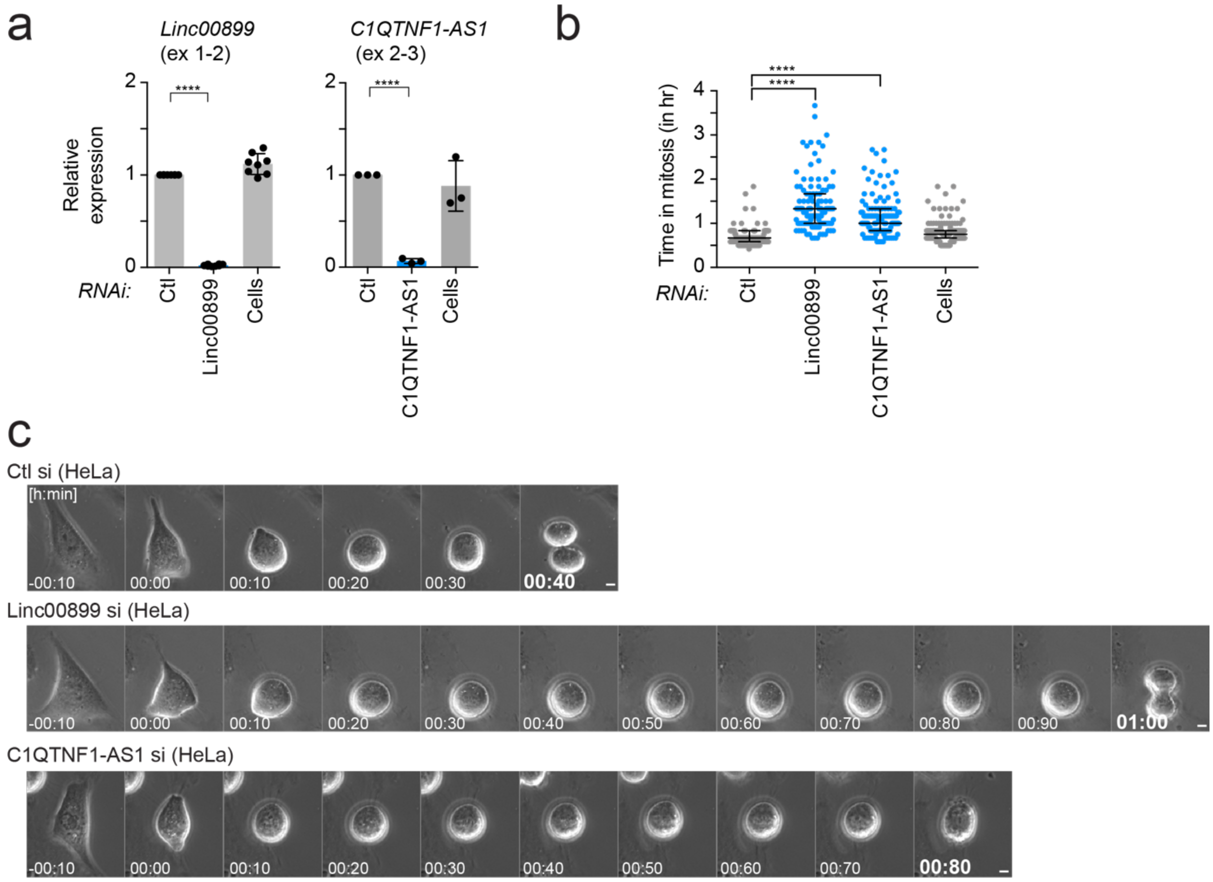


**Supplementary Fig. 3. RNAi-mediated depletion of *linc00899* and *C1QTNF1-AS1* leads to mitotic delay in HeLa cells.**

***a.*** Expression of *linc00899* and *C1QTNF1-AS1* after RNAi-mediated depletion of each lncRNA using pool of four siRNA sequences in HeLa cells, as quantified by qPCR. Results are also shown for cells treated with negative control siRNA (Ctl, from Ambion) and cells treated with transfection reagent alone (Cells). Expression values were compared to cells treated with negative control siRNA. Error bars, mean ± S.E.M. n = 3-8 biological replicates. Statistical significance by two-tailed Student's t -test: **** P<0.0001.

***b.*** Quantification of mitotic progression by time-lapse microscopy imaging of HeLa cells after RNAi-mediated depletion of *linc00899* and *C1QTNF1-AS1*. Mitotic duration was defined from nuclear envelope breakdown (NEBD, t=0 mins) to anaphase onset using bright-field microscopy. Number of cells analysed is n=126 for negative control siRNAs (Ctl), n=118 for *linc00899* RNAi, n=129 for *C1QTNF1-AS1* RNAi and n=140 for cells treated with transfection reagent alone (Cells). Bars show the median with interquartile range from 2 biological replicates. Statistical significance by Mann-Whitney test: ****P <0.0001.

***c.*** Representative still images from time-lapse microscopy imaging show the mitotic delay in *linc00899*- and *C1QTNF1-AS1*-depleted HeLa cells compared to cells treated with negative control siRNAs (Ctl). Scale bar, 5µm.

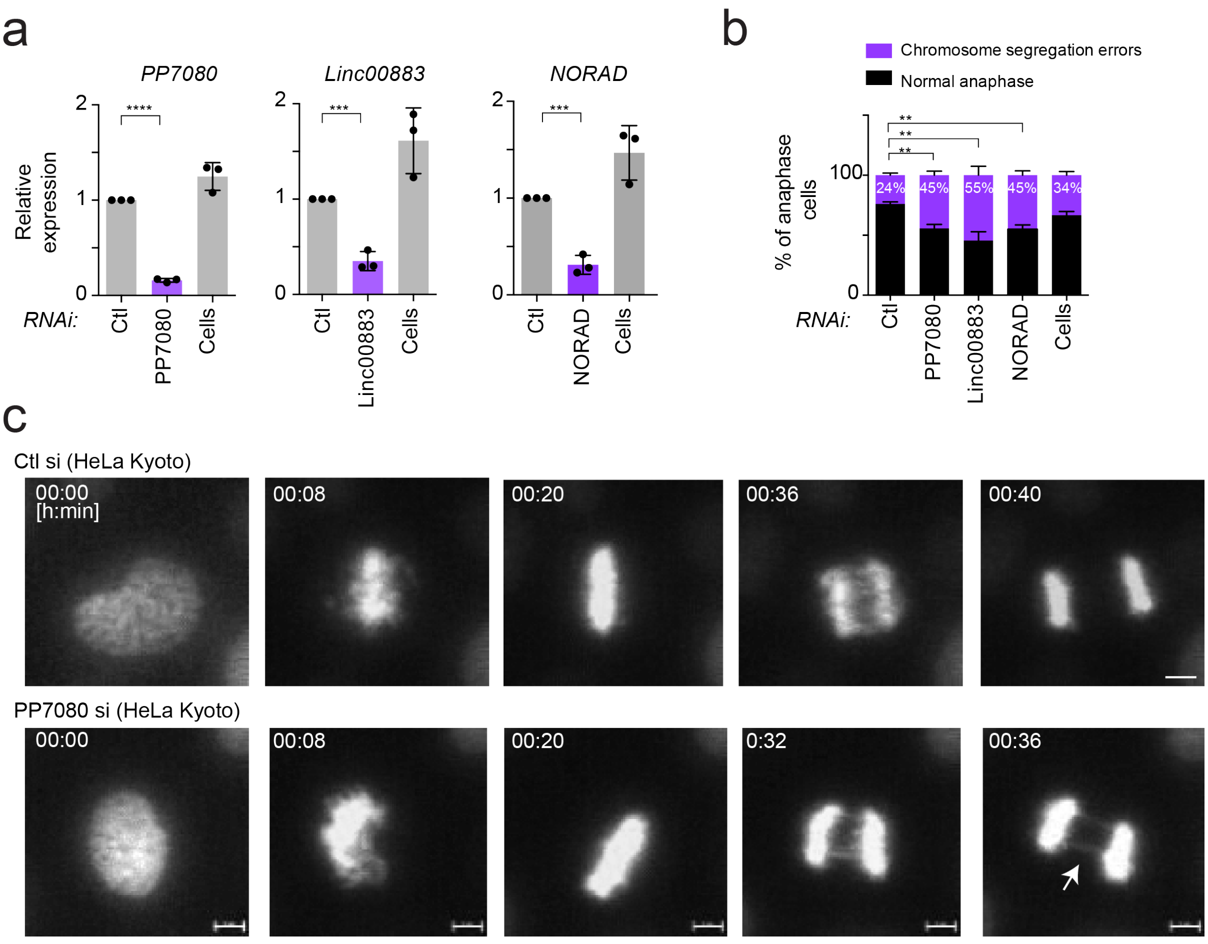


**Supplementary Fig. 4. RNAi-mediated depletion of *PP7080, linc00883* and *NORAD* leads to increased rate of chromosome segregation errors in HeLa cells.**

***a.*** Expression of *PP7080 (ENSG00000188242), linc00883 (ENSG00000243701) and NORAD* after RNAi-mediated depletion of each lncRNA using pool of four siRNA sequences in HeLa cells, as measured by qPCR. Results are also shown for cells treated with negative control siRNA (Ctl, from Ambion) and cells treated with transfection reagent alone (Cells). Expression levels were compared to the cells treated with control siRNA. Error bars represent mean ± S.E.M. n = 3 biological replicates. Statistical significance by two-tailed Student's t -test: *** P<0.001 and **** P<0.0001.

***b.*** Quantification of chromosome segregation errors in HeLa Kyoto cells after depletion of each lncRNA with RNAi using time-lapse microscopy imaging.

Number of anaphase cells analysed is n=110 for cells treated with transfection reagent alone (Cells), n=164 for negative control siRNAs (Ctl), n=140 for *NORAD* RNAi, n=140 for *PP7080* RNAi, and n=171 for *linc00883* RNAi. Each bar represents the mean of four biological replicates ± S.E.M. Statistical significance by two-tailed Student's t-test, **P <0.01.

***c.*** Representative still images of H2B-mCherry (white) from the time-lapse microscopy imaging in *PP7080*-depleted HeLa Kyoto cells expressing compared to cells treated with negative control siRNAs (Ctl). White arrow depicts chromosome segregation errors (chromatin bridges). Scale bar, 5µm.

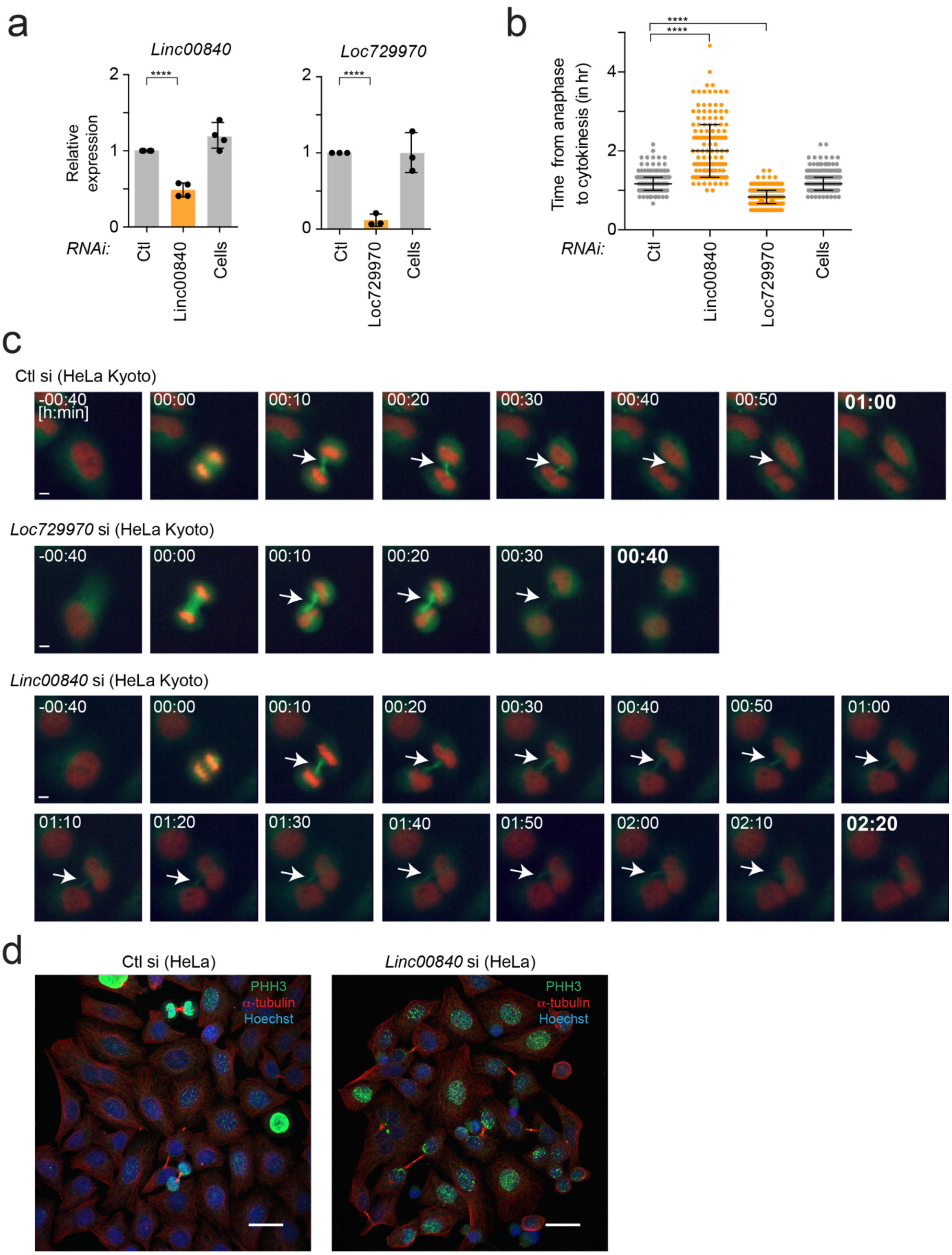


**Supplementary Fig. 5. RNAi-mediated depletion of *linc00840* and *loc729970* leads to cytokinesis defects in HeLa cells.**

***a.*** Expression of *linc00840 (ENSG00000226808)* and *loc729970 (ENSG00000235501)* after RNAi-mediated depletion of each lncRNA using a pool of four siRNA sequences in HeLa cells, as measured by qPCR. Results are also shown for cells treated with negative control siRNAs (Ctl, from Ambion) or cells treated with transfection reagent alone (Cells). Error bars, S.E.M. n = 3 - 4 biological replicates. Statistical significance by two-tailed Student's t -test: **** $P<0.0001$.

***b.*** Quantification of cytokinesis defects in HeLa Kyoto cells expressing eGFP $\alpha$-tubulin (green) and H2B-mCherry (red) and progressing through mitosis after depletion of each lncRNA with RNAi, as measured by time-lapse microscopy imaging. The time in cytokinesis was measured from anaphase onset (t=0 mins) to completion of abscission. The number of cells analysed was n=160 for cells treated with negative control siRNAs (Ctl), n=187 for cells treated with transfection reagent alone (Cells), n=130 for *linc00840* RNAi and n=238 for *loc729970* RNAi. Bars show the median with interquartile range from 3 biological replicates. Statistical significance by Mann-Whitney test: ****$P <0.0001$.

***c.*** Representative still images from the time-lapse microscopy in *linc00840* and *loc729970*-depleted HeLa Kyoto cells showing the time required to cleave the cytokinetic bridge, compared to cells treated with negative control siRNAs (Ctl). White arrow depicts cytokinetic bridge. Scale bar, 5µm.

***d.*** Representative confocal images of control and *linc00840*-depleted HeLa cells stained with $\alpha$-tubulin, PHH3 and Hoechst (DNA). Scale bar, 20µm. Note increased number of cytokinetic bridges in *linc00840*-depleted HeLa cells.

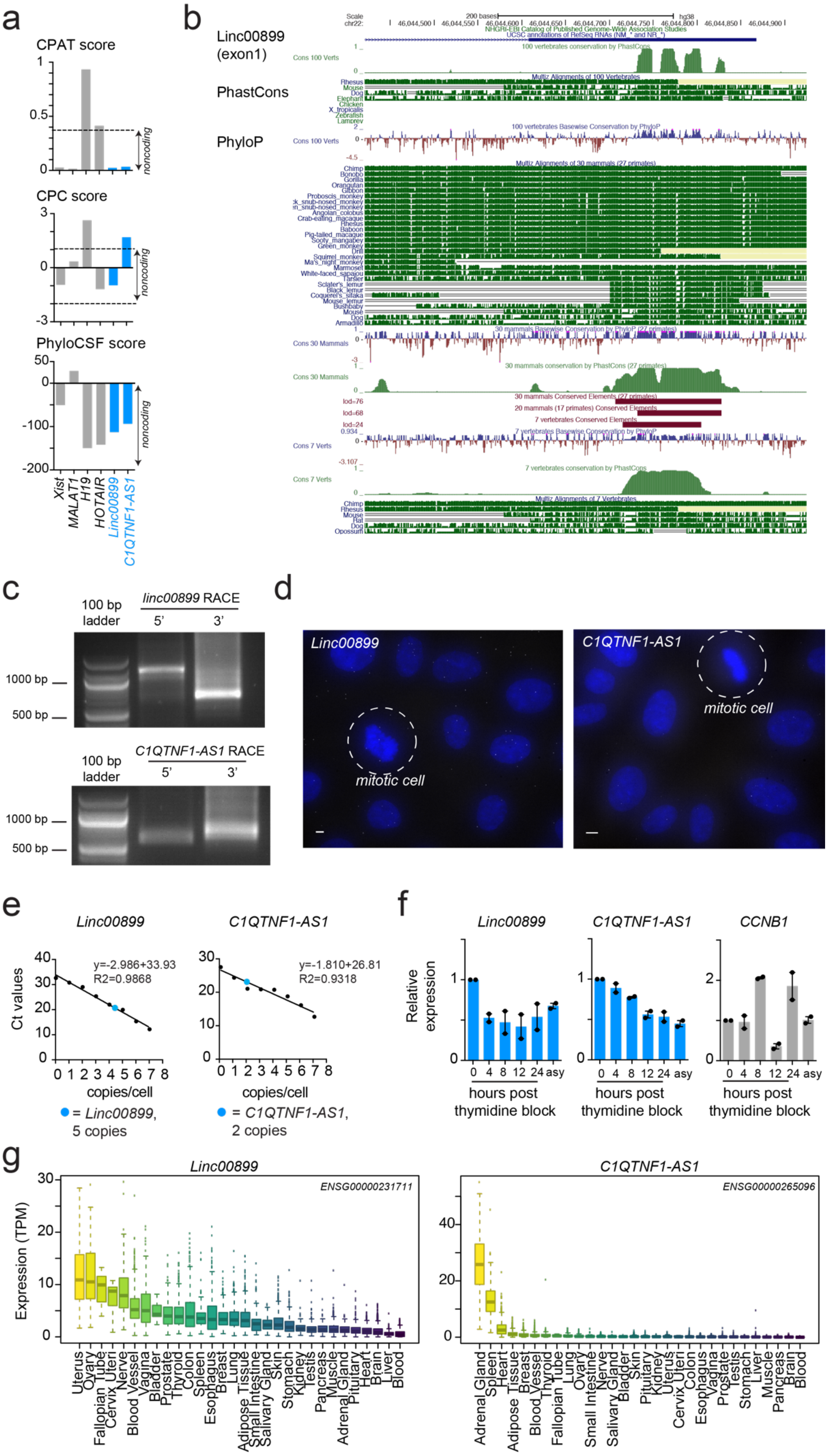

a
CPAT score
1
0.5
0
noncoding
CPC score
3
0
-3
noncoding
PhyloCSF score
50
0
-100
-200
noncoding
Xist
MALAT1
H19
HOTAIR
Linc00899
C1QTNF1-AS1
b
Linc00899 (exon1)
PhastCons
PhyloP
c
100 bp ladder
linc00899 RACE
5'
3'
1000 bp
500 bp
100 bp ladder
C1QTNF1-AS1 RACE
5'
3'
1000 bp
500 bp
d
Linc00899
mitotic cell
C1QTNF1-AS1
mitotic cell
e
Linc00899
y=-2.986+33.93
R2=0.9868
Ct values
copies/cell
= Linc00899, 5 copies
C1QTNF1-AS1
y=-1.810+26.81
R2=0.9318
copies/cell
= C1QTNF1-AS1, 2 copies
f
Linc00899
C1QTNF1-AS1
CCNB1
Relative expression
0 4 8 12 24 asy
hours post thymidine block
g
Linc00899
ENSG00000231711
Expression (TPM)
Uterus
Ovary
Fallopian Tube
Cervix Uteri
Nerve
Blood Vessel
Vagina
Bladder
Prostate
Thyroid
Colon
Spleen
Esophagus
Breast
Lung
Adipose Tissue
Small Intestine
Salivary Gland
Skin
Stomach
Kidney
Testis
Pancreas
Muscle
Adrenal Gland
Pituitary
Heart
Brain
Liver
Blood
C1QTNF1-AS1
ENSG00000265096
Adrenal Gland
Spleen
Heart
Adipose Tissue
Breast
Blood Vessel
Thyroid
Fallopian Tube
Lung
Ovary
Small Intestine
Nerve
Salivary Gland
Bladder
Skin
Pituitary
Kidney
Uterus
Cervix Uteri
Colon
Esophagus
Vagina
Prostate
Testis
Stomach
Liver
Muscle
Pancreas
Brain
Blood

**Supplementary Fig. 6. Molecular and cellular characteristics of *linc00899* and *C1QTNF1-AS1.***

***a.*** Computational analysis of the *linc00899* and *C1QTNF1-AS1* sequences using a variety of protein coding potential tools. Low protein-coding potentials were detected by CPAT, CPC and PhyloCSF score. For comparison, scores are also shown for a number of well-known lncRNAs as positive controls (black text).

***b.*** Sequence conservation of exon 1 of *linc00899,* as scored with PhastCons and PhyloP. Results are shown as tracks on the UCSC genome browser.

***c.*** Identification of the 5' and 3' ends of *linc00899* and *C1QTNF1-AS1* with RACE in HeLa cells. Isoforms were identified with lengths ranging from 1144 to 1562 bp for *linc00899* and 864 to 952 bp for *C1QTNF1-AS1*.

***d.*** Single molecule RNA FISH of *linc00899* (left) and *C1QTNF1-AS1* (right) in interphase and mitotic cells (circle) using exonic probes against each of the mature transcript. Representative images are shown from at least 3 biological replicates. DNA is stained with DAPI and shown in blue, while probes are shown in white. Scale bar represents 20µm.

***e.*** Copy numbers of *linc00899* and *C1QTNF1-AS1* in HeLa cells, determined by qPCR. RNA was extracted from known number of HeLa cells, and the copy number per cell was computed using the standard curve method for Ct values against dilutions of a lncRNA DNA template of known concentration. *Linc00899* is present at five copies while *C1QTNF1-AS1* at two copies per cell. n = 5 biological replicates.

***f.*** Expression levels of *linc00899* and *C1QTNF1-AS1* in synchronized HeLa cells using double thymidine block as measured by qPCR. *Cyclin B1 (CCNB1)* was used as a positive control to mark cells in G2/M phase of the cell cycle.

Expression levels were standardised to time point 0 hr. Results are also shown for asynchronous cells (asy). Error bars, S.E.M. n = 2 biological replicates.

***g.*** Expression values for *linc00899* and *C1QTNF1-AS1* were obtained from version 7 of the GTEx RNA-seq expression dataset (*https://gtexportal.org/home/datasets*) for normal tissues and are shown as boxplots of transcripts per million (TPM). Tissues are sorted in order of decreasing median expression across all GTEx samples for each lncRNA. Points for each tissue represent samples that lie more than 1.5-fold interquartile ranges from the third quartile.

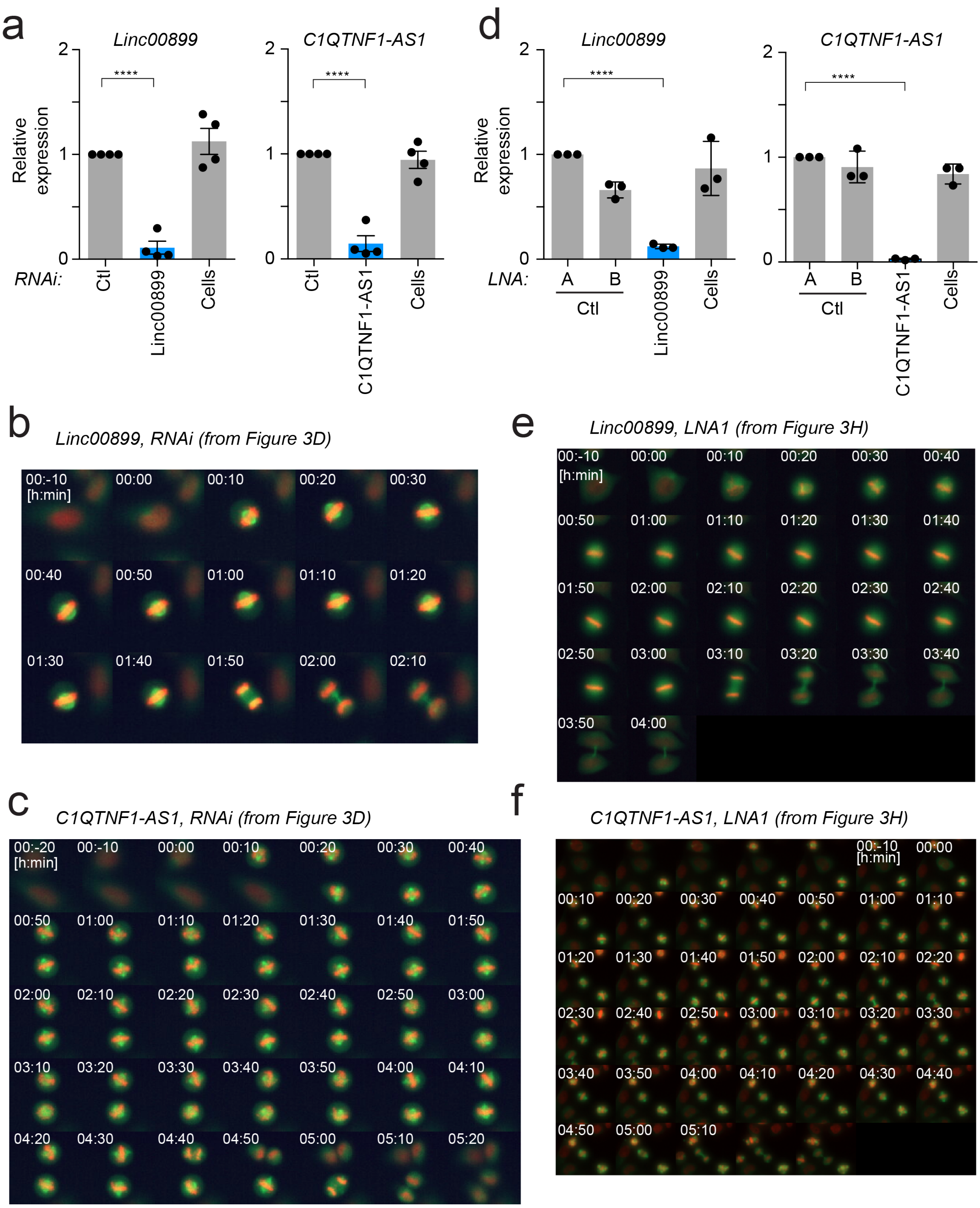


**Supplementary Fig. 7. RNAi- and LNA-mediated depletion of *linc00899* and *C1QTNF1-AS1* in HeLa Kyoto cells.**

***a.*** Expression of *linc00899 and C1QTNF1-AS1* in HeLa Kyoto cells after their depletion with RNAi using a pool of four siRNA sequences, as measured with qPCR. Results are also shown for cells treated with negative control siRNAs (Ctl, from Ambion) or cells treated with transfection reagent alone (Cells). Error bars, S.E.M. n = 4 biological replicates. Statistical significance by two-tailed Student's t -test: **** P<0.0001.

***b.*** One representative series of still images from time-lapse microscopy in *linc00899*-depleted HeLa Kyoto cells.

***c.*** One representative series of images from time-lapse microscopy in *C1QTNF1-AS1*-depleted HeLa Kyoto cells.

***d.*** Expression of *linc00899* and *C1QTNF1-AS1* in HeLa Kyoto cells after their depletion with LNA1 gapmers, as measured with qPCR. Results are also shown for cells treated with negative control LNA A and B (Ctl A and B) or with transfection reagent alone (Cells). Error bars, S.E.M. n = 3 biological replicates. Statistical significance by two-tailed Student's t -test: **** P<0.0001.

***e.*** One representative series of still images from time-lapse microscopy in *linc00899*-depleted HeLa Kyoto cells.

***f.*** One representative series of images from time-lapse microscopy in *C1QTNF1-AS1*-depleted HeLa Kyoto cells.

The cropped series are shown in Figure 3.

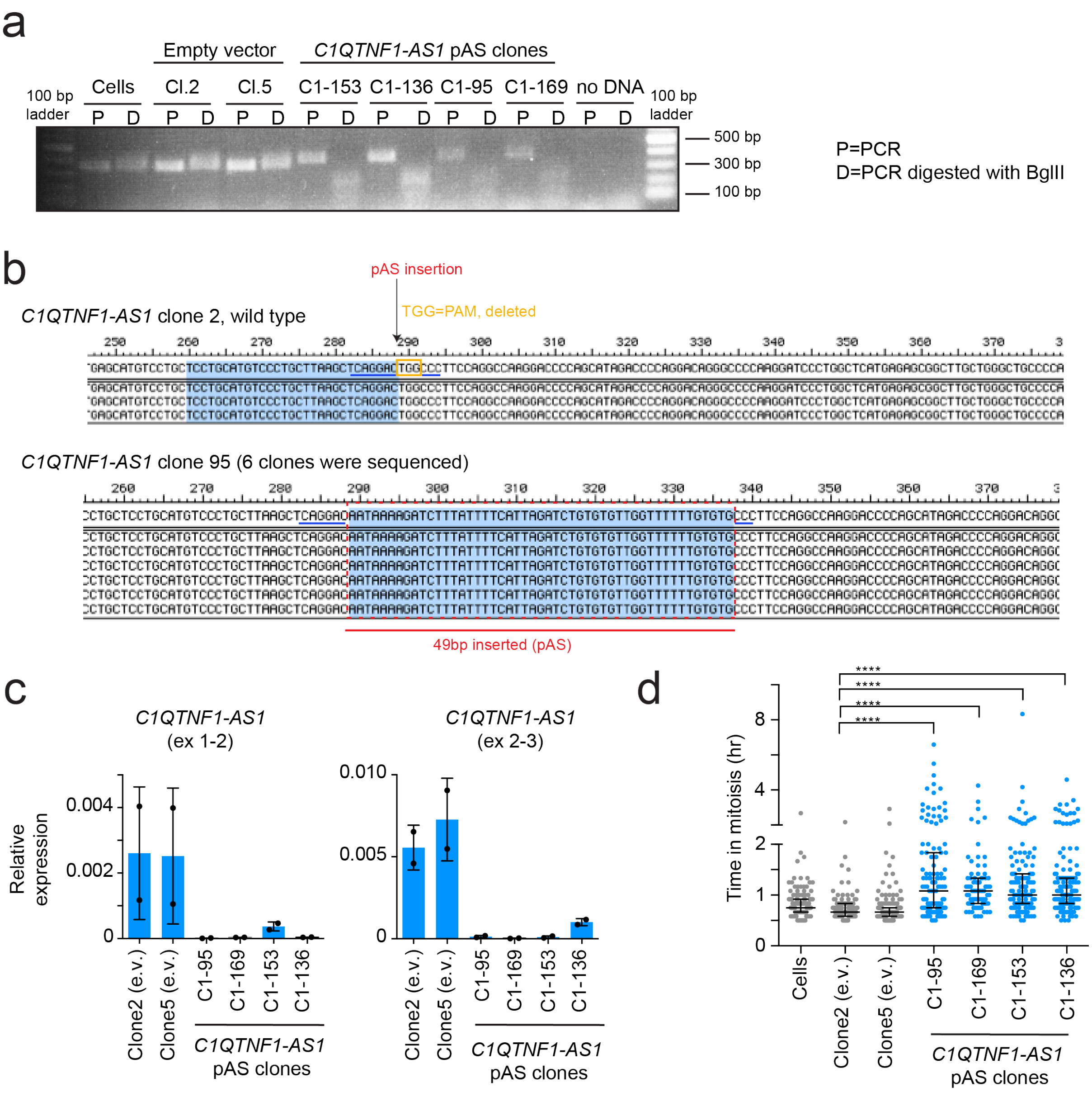


**Supplementary Fig. 8. Generation of CRISPR clones with insertion of poly(A) signal to terminate transcription of *C1QTNF1-AS1.***

***a.*** Detection of genomic DNA containing poly(A) sequence (pAS) insertion at the *C1QTNF1*-AS1 locus, as determined by PCR. Results are shown for HeLa cells alone (Cells), clones stably integrated with an empty vector PX458 (e.v., clones 2 and 5, negative controls), and four different *C1QTNF1-AS1* clones transfected with the pAS-*C1QTNF1-AS1* insert (clones C1-95, C1-169, C1-153 and C1-136). After PCR, a small aliquot was digested with *Bgl*II. In cells with no insert, the

PCR product should be 358 bp and should not be affected by digestion. If pAS (49 bp) was inserted, the PCR product should be 404 bp (49 bp+358 bp) before digestion, and should yield two products of 149 bp and 237 bp after digestion. The 100 bp ladder is shown on each side of the 2% agarose gel. PCR primers are present in the Supplementary Information.

***b.*** Representative sequences after pAS insertion into the *C1QTNF1-AS1* sequence from clone 95, compared to negative control clone 2. Genomic DNA was extracted from the cells and cloned into pJET BLUNT. After transformation, several bacteria clones were subjected to Sanger sequencing using pJET Forward primers (Supplementary Information).

***c.*** Expression of *C1QTNF1-AS1* after insertion of pAS into exon 1 of the *C1QTNF1-AS1* locus, as quantified by qPCR. The pAS insertion allows transcription at the *C1QTNF1-AS1* locus while inhibiting production of the *C1QTNF1-AS1* transcript. Results are shown for cells transfected with the empty vector PX458 (e.v., clones 2 and 5 as negative controls) and four pAS-*C1QTNF1-AS1* clones. qPCR primers against exons 1-2 and 2-3 were used to quantify the knockdown efficiency of *C1QTNF1-AS1*. Error bars, S.E.M. n = 2 biological replicates.

***d.*** Quantification of mitotic progression by time-lapse microscopy imaging in four HeLa *C1QTNF1-AS1* pAS clones. Mitotic duration was measured from nuclear envelope breakdown (NEBD, t=0 mins) to anaphase onset using the bright-field microscopy. The number of mitotic cells analysed was n=137 for HeLa cells alone (Cells), n=109 for the negative control clone 2, n=145 for negative control clone 5, n=121 for *C1QTNF1-AS1* pAS clone 95, n=84 for pAS clone 169, n=131 for pAS clone 153, and n=156 for pAS clone 136. Bars show median with interquartile range from two biological replicates. Statistical significance by Mann-Whitney test: ****P <0.0001.

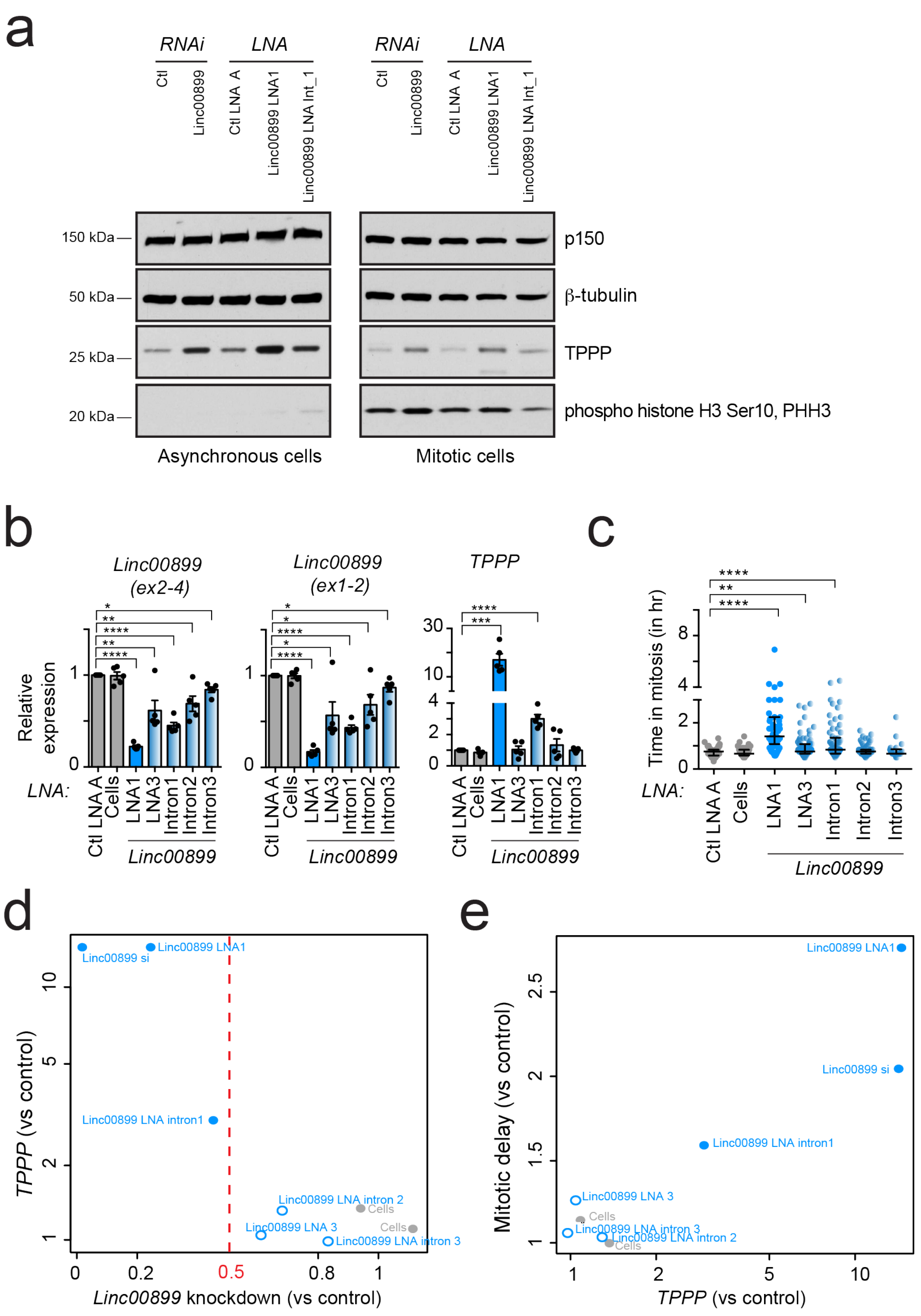


**Supplementary Fig. 9. Depletion of *linc00899* using additional LNA gapmers targeting either exon 1 (LNA 3) or intronic regions of *linc00899* (intronic LNAs).**

***a.*** Western blot of TPPP levels after depletion of *linc00899* in asynchronous (left) and mitotic cells (right) compared to negative control siRNA (Ctl, from Ambion) or negative control LNA (Ctl LNA A). β-tubulin and p150 were used as loading controls. Mitotic cell extracts were obtained by mitotic shake-off of monastrol-treated cells and PHH3 antibody was used as a control to show enrichment of mitotic cells.

***b.*** Expression of *linc00899* and *TPPP* after depletion of *linc00899* using LNA gapmer 1 (as in Figure 3) with additional LNA gapmers targeting either different region of exon 1 (LNA 3) or different intronic regions (intron 1-3). Results are also shown for cells treated with negative control LNA A (Ctl LNA A) and for cells treated with transfection reagent alone (Cells). Expression values are shown relative to cells treated with negative control LNA A. Error bars, S.E.M. n = 5 biological replicates. Statistical significance by two-tailed Student's *t*-test: * P<0.1, ** P<0.01, *** P<0.001 and **** P<0.0001.

***c.*** Quantification of mitotic progression in HeLa cells by time-lapse microscopy imaging after LNA-mediated depletion of *linc00899* using the additional LNA gapmers as in (***b***). Mitotic duration was measured from NEBD (t=0 mins) to anaphase onset by bright-field microscopy. Number of mitotic cells analyzed was n=97 for negative control LNA A (Ctl LNA A), n=51 for cells treated with transfection reagent alone (Cells), n=54 for *linc00899* LNA 1, n=83 for *linc00899* LNA 3, n=75 for *linc00899* LNA targeting intron 1, n=79 for *linc00899* LNA targeting intron 2, and n=23 for *linc00899* LNA targeting intron 3. Bars show median with interquartile range from 2 biological replicates. Statistical significance by Mann-Whitney test: ***P*<0.01 and *****P*<0.0001.

***d.*** Upregulation of *TPPP* expression with respect to *linc00899* knockdown efficiency, after depleting *linc00899* with a variety of LOF methods. Expression was quantified using qPCR and calculated relative to the appropriate negative control for each method. Each value represents the average of 4-5 replicates per condition. Results are also shown for cells treated with transfection reagent alone (Cells, in grey). LOF methods achieving more than 50% depletion are denoted with the blue closed circles.

***e.*** Fold increase in the mitotic delay upon depletion of *linc00899* with a variety of LOF methods, with respect to *TPPP* upregulation. All values were computed relative to the appropriate negative control for each LOF method and represent the average of 4-5 replicates per condition. Points are denoted as described in (***b***).

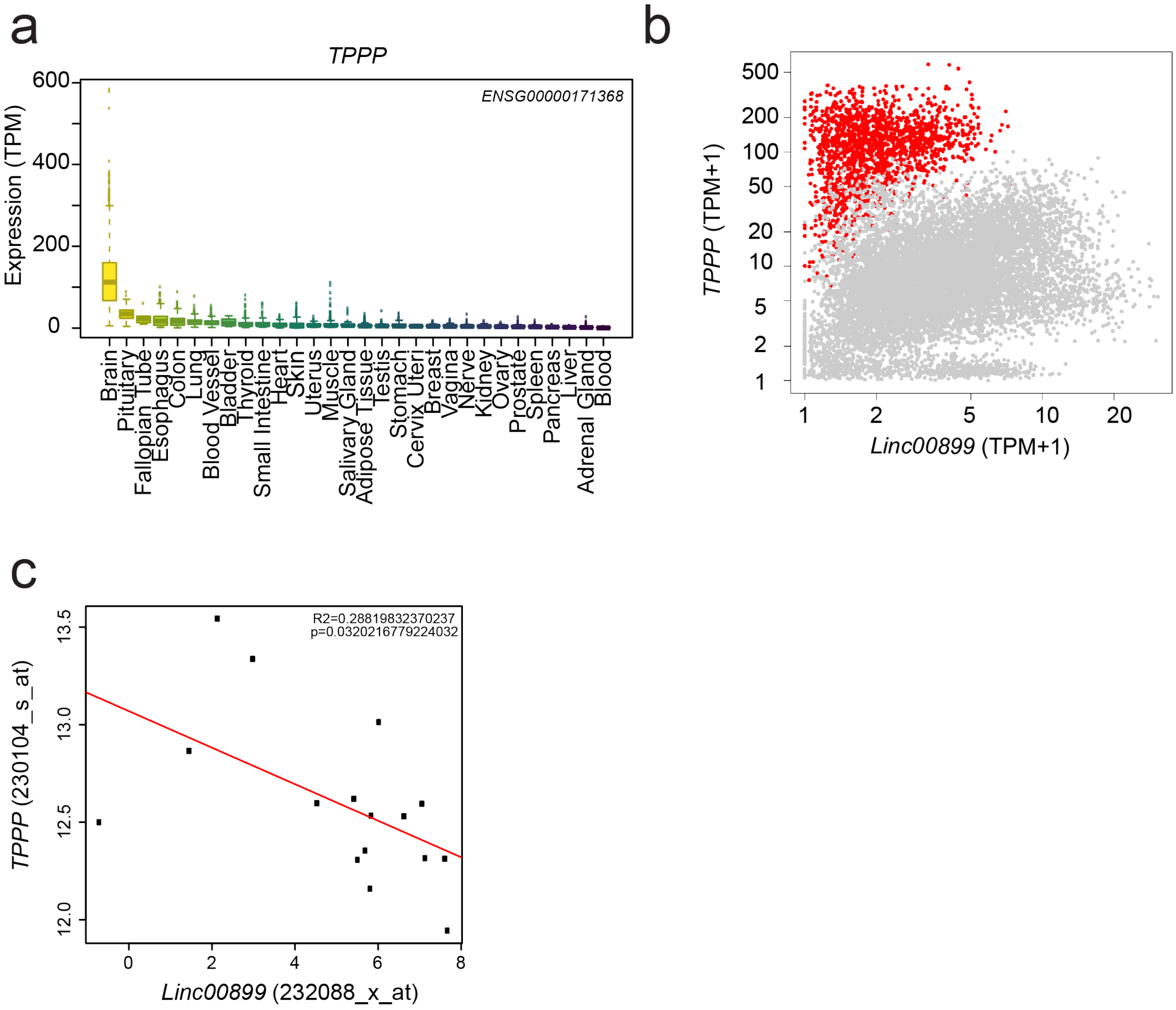


**Supplementary Fig. 10. Expression of *TPPP* and *linc00899* in humans.**

***a.*** Boxplots of *TPPP* expression in transcripts-per-million (TPM) across all samples from each tissue. Tissues are sorted by decreasing median expression.

***b.*** Expression of *TPPP* relative to *linc00899* across all samples in the GTEx dataset. Each point represents a single sample, with all brain samples coloured in red.

***c.*** Expression of *TPPP* relative to *linc00899* across all samples in the GSE52139 dataset. Each point corresponds to one multiple sclerosis patient, and all gene expression values represent $log_2$-transformed probe intensities. The line of best fit is shown in red, with an $R^2$ value of 0.29 and a p-value of 0.03.

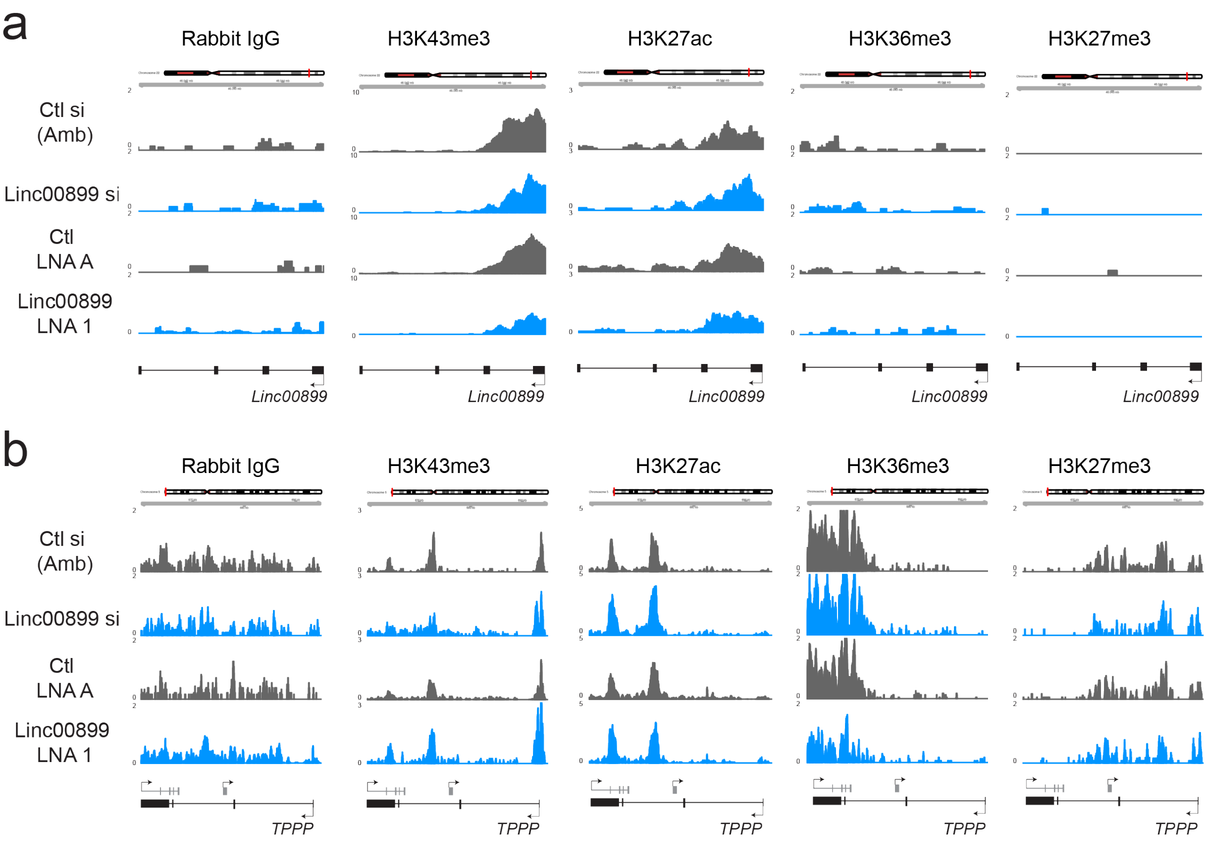


**Supplementary Fig. 11. Chromatin landscape at the *linc00899* and *TPPP* locus after depletion of *linc00899*.**

CUT&RUN profiling of active (H3K4me3, H3K27ac, H3K36me3) and repressive (H3K27me3) histone modifications at the *linc00899* (***a***) and *TPPP* (***b***) locus after depletion of *linc00899* with RNAi and LNAs in HeLa cells. Coverage tracks are also shown for negative control siRNAs (Ctl, from Ambion) and negative control LNA A (Ctl LNA A). Each track represents the average normalized count-per-million at each base position across two biological replicates.

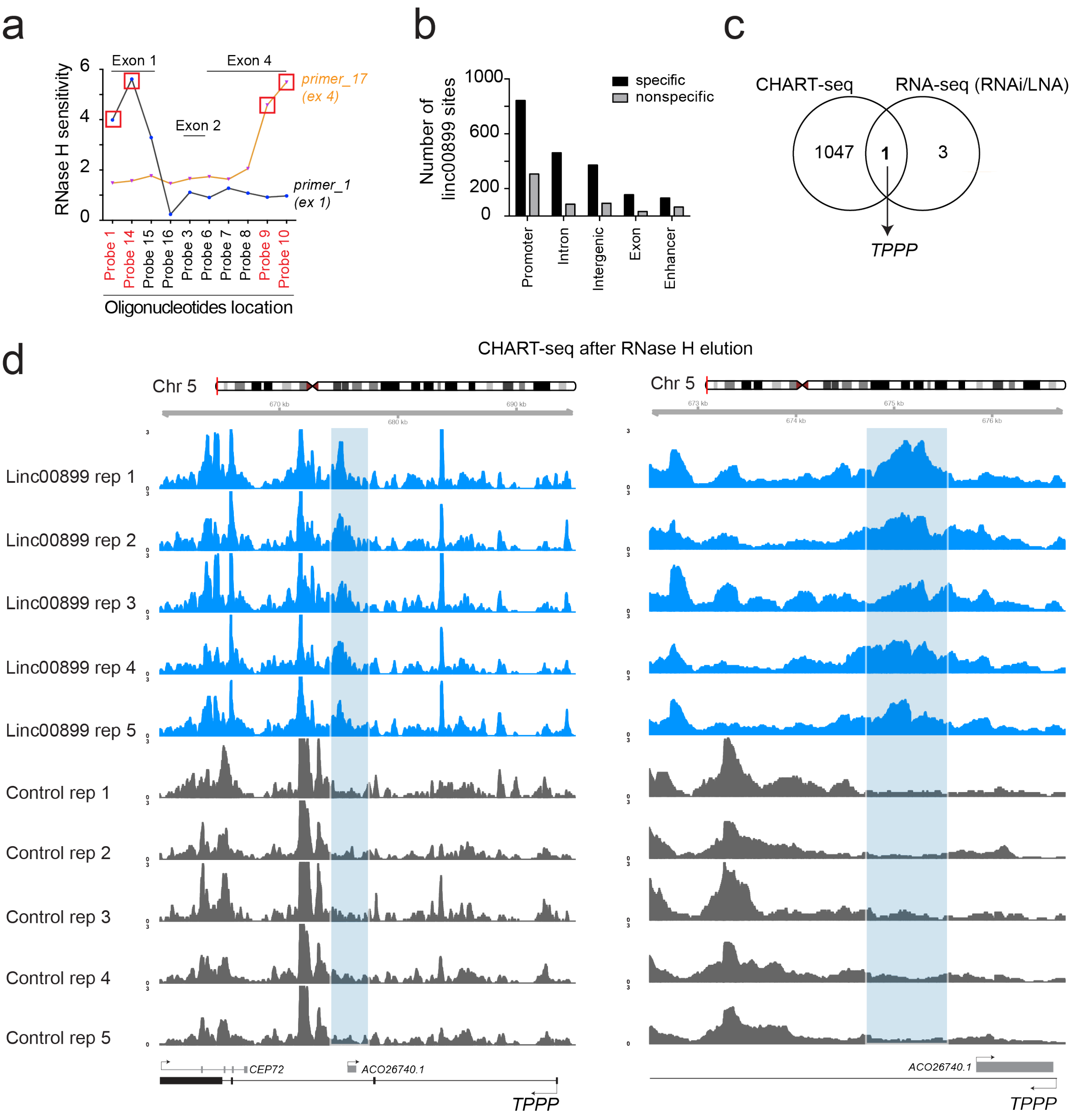


**Supplementary Fig. 12. Statistics for *linc00899* CHART-seq.**

***a.*** Mapping oligonucleotide binding to the *linc00899* transcript by RNase H sensitivity assays. Several antisense oligonucleotides were designed that directly hybridized to endogenous *linc00899* and RNase H mapping was used to identify oligonucleotides binding to accessible regions of the transcript. A cocktail of four such oligonucleotides were used to purify chromatin complexes containing *linc00899* transcript, whereas sense DNA oligonucleotides were used as controls to account for any non-specific binding of the probes to the DNA locus. Each DNA oligonucleotide was added to cross-linked, sheared chromatin from HeLa cells to form RNA-DNA hybrids with the lncRNA transcript. Samples were treated with RNase H to digest the RNA in the RNA-DNA hybrids and incubated with DNase I to remove genomic DNA. RT-qPCR was performed to quantify RNase H cleavage of *linc00899*, using primers targeting exon 1 (primer 1) or exon 4 (primer 17) of *linc00899*. RNase H sensitivity was calculated based on the ratio of cleaved to uncleaved transcript, using no oligonucleotide as a control reaction. Different oligonucleotides spanning the *linc00899* transcript were tested, and the cocktail mix of probe 1 (exon 1), probe 14 (exon 1), probe 9 (exon 4) and probe 10 (exon 4) was used for CHART-seq.

***b.*** Number of *linc00899* binding sites in different genomic contexts at an empirical FDR of 30%. We identified ~1964 locations with significant increases in coverage upon antisense pulldown compared to sense control. These putative *linc00899* binding sites were mostly distributed within promoters and introns. Specific binding sites were defined as those with significantly increased coverage in the antisense pulldown, while non-specific sites were defined as those with increased coverage in the sense pulldown (i.e., the negative control).

***c.*** Overlap between the set of genes (1048) bound by *linc00899* in CHART-seq and the set of genes that were differentially expressed in the same direction in RNA-seq after LNA or RNAi-mediated depletion (see Fig. 4b). The only gene in the intersection was *TPPP*.

***d.*** CHART-seq coverage tracks at the *TPPP* locus after pulldown with antisense (blue) or sense oligonucleotides (grey). Coverage is shown for 5 biological replicates (left), with a zoomed-in view of the site with the most significant change in coverage (right).

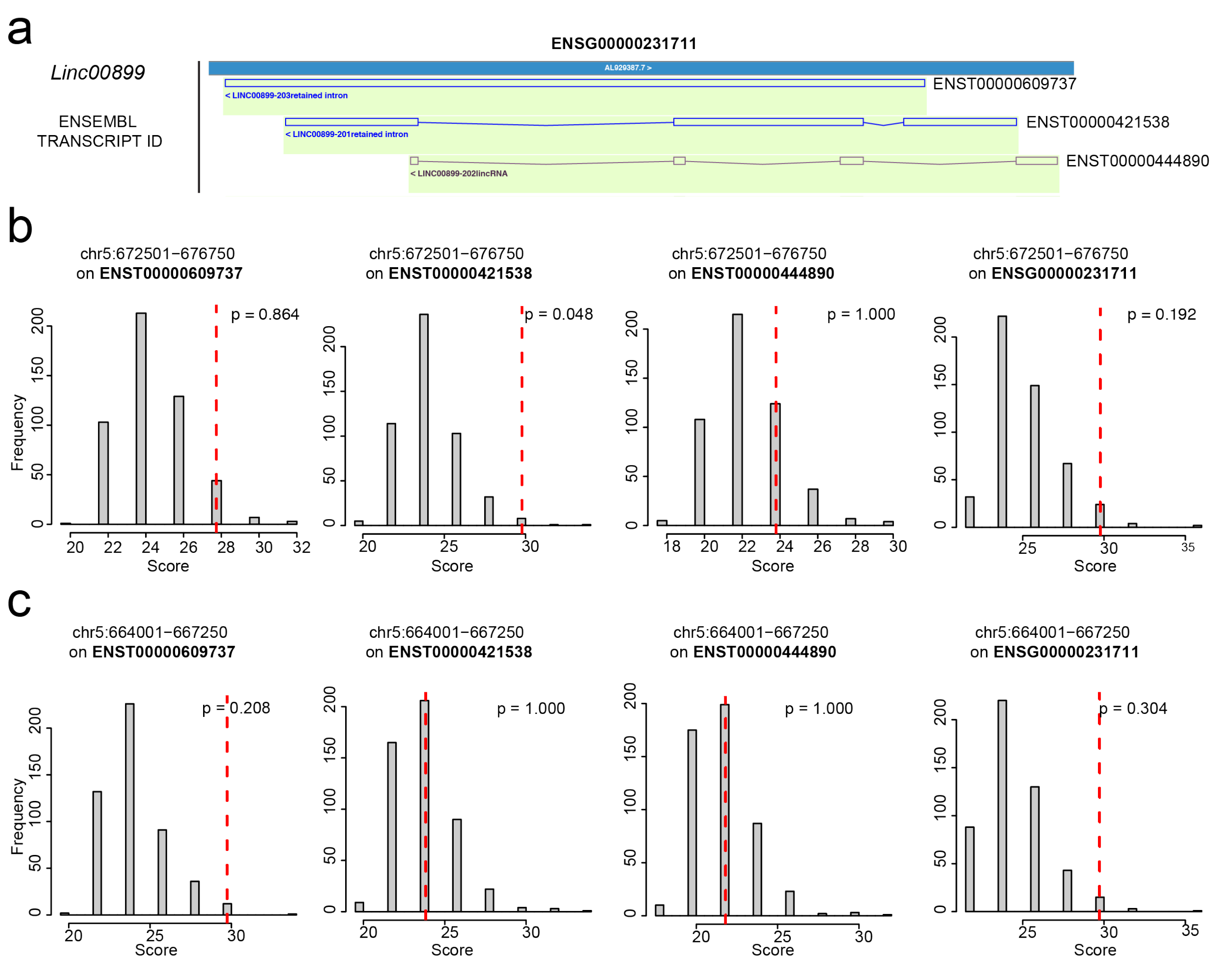


**Supplementary Fig. 13. Complementarity between the *TPPP* genomic sequence and the transcript sequence of *linc00899.***

Smith-Waterman local alignment scores between the *linc00899* transcript and the sequences of the genomic intervals within *TPPP* bound by *linc00899* (as detected by CHART-seq). We examined three isoforms of *linc00899* from Ensembl annotation along with the premature transcript (***a***). For each interval (***b, c***), we performed a local alignment between each *linc00899* sequence to the genome sequence, taking the maximum score across both strands of the latter. We repeated this process after shuffling the sequence to obtain a null distribution of 100 alignment scores. Genuine matches should exhibit a real alignment score (red) that is much greater than the random distribution (grey).

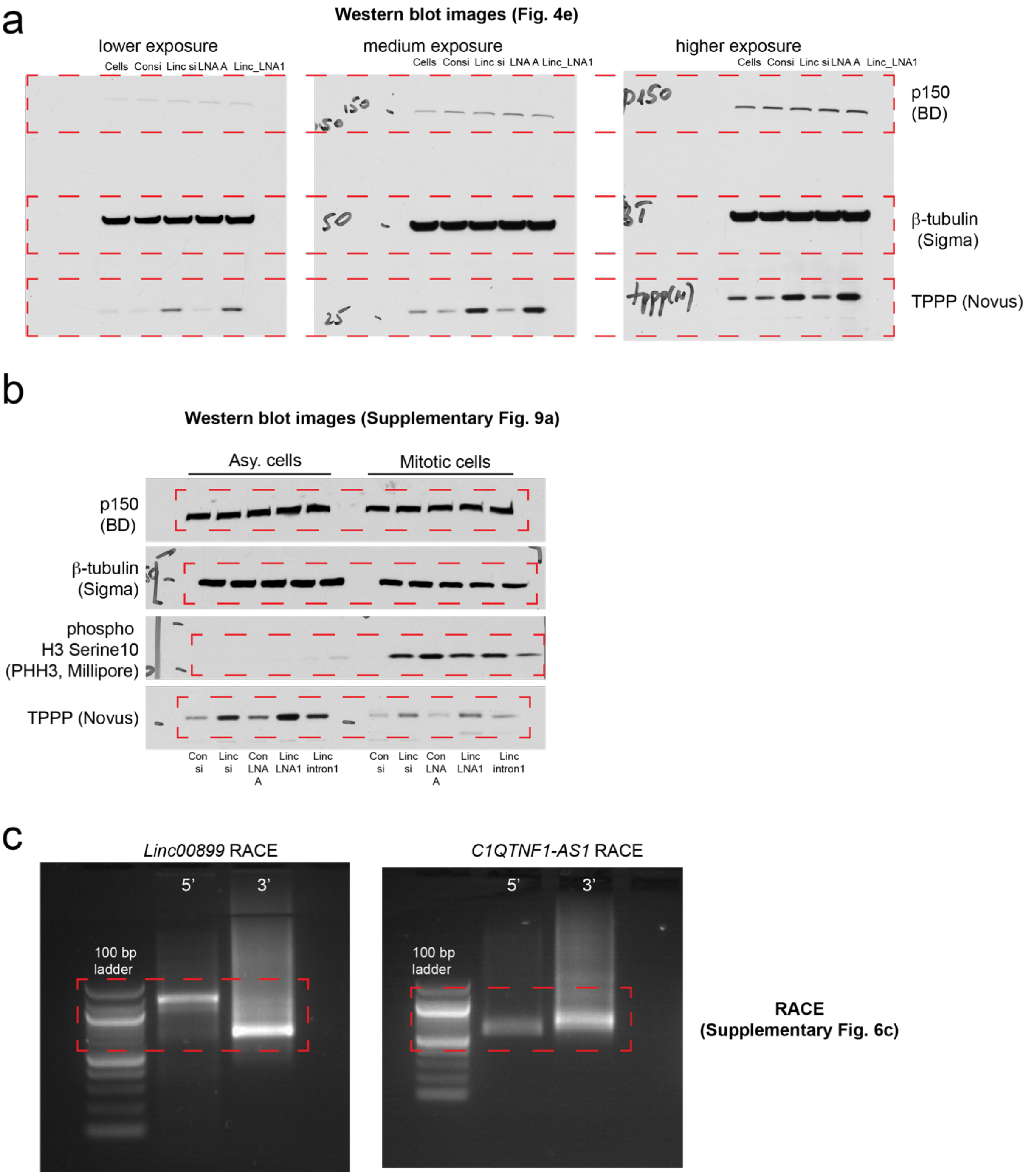


**Supplementary Fig. 14. Uncropped images of the western blots and gels.** Regions in red boxes are shown in the main text for Fig. 4e (***a***), Supplementary Fig. 9a (***b***) and Supplementary Fig. 6c (***c***). The RACE products were analysed on a 1% agarose gel.
